# A conserved role of the insulin-like signaling pathway in uric acid pathologies revealed in *Drosophila melanogaster*

**DOI:** 10.1101/387779

**Authors:** Sven Lang, Neelanjan Bose, Kenneth A. Wilson, Deanna J. Brackman, Tyler Hilsabeck, Mark Watson, Jennifer N. Beck, Amit Sharma, Ling Chen, David W. Killilea, Sunita Ho, Arnold Kahn, Kathleen Giacomini, Marshall L. Stoller, Thomas Chi, Pankaj Kapahi

**Affiliations:** The Buck Institute for Research on Aging, Novato, California, 94945, United States of America.; Department of Bioengineering and Therapeutic Sciences, University of California San Francisco, San Francisco, California, 94158, United States of America.; Division of Biomaterials and Bioengineering, University of California San Francisco, San Francisco, California, 94143, United States of America.; Nutrition & Metabolism Center, Children’s Hospital Oakland Research Institute, Oakland, California, 94609, United States of America.; Department of Urology, University of California San Francisco, San Francisco, California, 94143, United States of America.; Davis School of Gerontology, University of Southern California, Los Angeles, California, 90089, United States of America.

**Keywords:** Uric acid, urate oxidase, gout, kidney stones, insulin-like signaling, FoxO, NADPH Oxidase, AKT, *Drosophila melanogaster*, purine

## Abstract

Elevated uric acid (UA) is a key factor for disorders, including gout or kidney stones and result from abrogated expression of *Urate Oxidase (Uro)* and diet. To understand the genetic pathways influencing UA metabolism we established a *Drosophila melanogaster* model with elevated UA using *Uro* knockdown. Reduced *Uro* expression resulted in the accumulation of UA concretions and diet-dependent shortening of lifespan. Inhibition of insulin-like signaling (ILS) pathway genes reduced UA and concretion load. In humans, SNPs in the ILS genes *AKT2* and *FOXO3* were associated with UA levels or gout, supporting a conserved role for ILS in modulating UA metabolism. Downstream of the ILS pathway UA pathogenicity was mediated partly by NADPH Oxidase, whose inhibition attenuated the reduced lifespan and concretion accumulation. Thus, genes in the ILS pathway represent potential therapeutic targets for treating UA associated pathologies, including gout and kidney stones.

**Highlights:**

- In *Drosophila* high uric acid (UA) levels shorten lifespan and cause UA aggregation
- Conserved in flies and humans, the ILS pathway associates with UA pathologies
- FoxO dampens concretion formation by reducing UA levels and ROS formation
- Inhibition of NOX alleviates the lifespan attenuation and UA aggregation

## Introduction

Elevated serum uric acid (UA) in humans often stems from the consumption of an enriched, so-called Westernized diet coupled with the lack of a functional urate oxidase gene (*Uro*). Such elevated UA levels in the blood (hyperuricemia) or urine (hyperuricosuria) are key risk factors for gout and kidney stones, respectively (Kuo et al., 2015; Zhu et al., 2012). Gout, which affects nearly 4% of the US population, often presents with acute arthritis that is caused by a buildup of UA crystals, typically in the metatarsal-phalangeal joint space (Fisher et al., 2014; Kuo et al., 2015). Patients with urinary stone disease often show elevated UA levels as well. More broadly, hyperuricemia affects over 20% of the US population and has been associated with hypertension, dyslipidemia, kidney disease, type 2 diabetes, premature death, and higher all-cause mortality risk (Barron et al., 2015; Fisher et al., 2017; Giordano et al., 2015; Kanbay et al., 2016; Kuwabara, 2016; Peng et al., 2015; Perez-Ruiz et al., 2015).

UA and its anion urate represent the final breakdown product of the human purine degradation pathway. Sequential mutational events in the promoter and coding sequence first reduced and then abolished expression of *Uro* during the evolution of the hominid lineage ~15 million years ago (Johnson et al., 2011; Oda et al., 2002). Hence, in contrast to many other animals humans are unable to enzymatically convert UA to the better water soluble product allantoin. While many mammals show serum UA levels of 1 mg/dl and lower, humans typically settle in a range of 4 - 6 mg/dl, close to the UA solubility limit of 6.8 mg/dl at physiological pH and body temperature (Grassi et al., 2013). Co-aligning with the expression pattern of the UA producing xanthine dehydrogenase the liver, as well as small and large intestine represent the main sites of UA production in the human body (Thul et al., 2017). While circulating UA is freely filtered at the renal glomerulus usually 95 - 99% of this fraction is reabsorbed in the proximal renal tubule via transporters such as URAT1 (SLC22A12) or GLUT9 (SLC2A9) and brought back into circulation (Enomoto et al., 2002; Vitart et al., 2008).

Despite the impact of elevated UA in several disease processes, there has been a relative lack of new therapeutics, in part because of a lack of good animal models through which to explore key mechanisms underlying UA accumulation and its sequelae. Consequently, we designed a *Drosophila melanogaster* model to study the impact of rising UA levels in a genetic and dietary tractable model organism recapitulating both aspects encountered in hominoid evolution: depletion of urate oxidase activity combined with the changes in dietary habits. This system paved the way to determine previously unaccounted, aging relevant modulators to reduce UA and associated pathologies.

## Results and Discussion

### *Uro* knockdown shortens lifespan and causes UA accumulation in a diet-dependent manner in *Drosophila*

Unlike humans, *Drosophila melanogaster* uses the enzyme urate oxidase to convert UA into allantoin (Fig. S1A). Therefore, we “humanized” a population of *Drosophila* to recapitulate the lack of functional urate oxidase making UA the end-product of purine catabolism. We used the heterologous, ligand (RU486)-induced gene switch system to generate spatially and temporally precise RNAi-mediated knockdown of *Uro* in *Drosophila* (McGuire et al., 2004). *Uro* knockdown was initiated in 2-3 day-old flies using the ubiquitous *daughterless* gene switch driver (Fig. S1B). Compared to isogenic control flies, *Uro* knockdown animals (Uro-RNAi #1) showed a diet-dependent shortening of lifespan (Fig. 1A). Specifically, Uro-RNAi flies fed diets containing yeast concentrations above 2% died sooner, on average. The peak effect was observed for those fed a 5%-containing high yeast (HY) diet (Fig. 1B). The diet-dependent lifespan attenuation was further confirmed using a second *Uro* targeting RNAi, and alternative ubiquitous (*Da-GAL4*) or Malpighian tubule-specific (*c42-GAL4*, *Uro-GAL4*) knockdown of *Uro* (Fig. S1B-S1F), with the latter being the main tissue expressing *Uro* (Chintapalli et al., 2007). As a negative control, knockdown of *Uro* using the neuronal *Elav-GAL4* driver, i.e., in a tissue devoid of *Uro* expression, failed to show any effect (Fig. S1E, S1F). Of note, consistent with the concept that caloric restriction can extend lifespan in various species (López-Otín et al., 2016; Soultoukis and Partridge, 2016) our control and knockdown flies are significantly longer lived on low nutrient diets with 0.5% and 1% yeast extract compared to diets with a higher yeast extract content (Fig. 1B, S1D).

**Figure 1.**
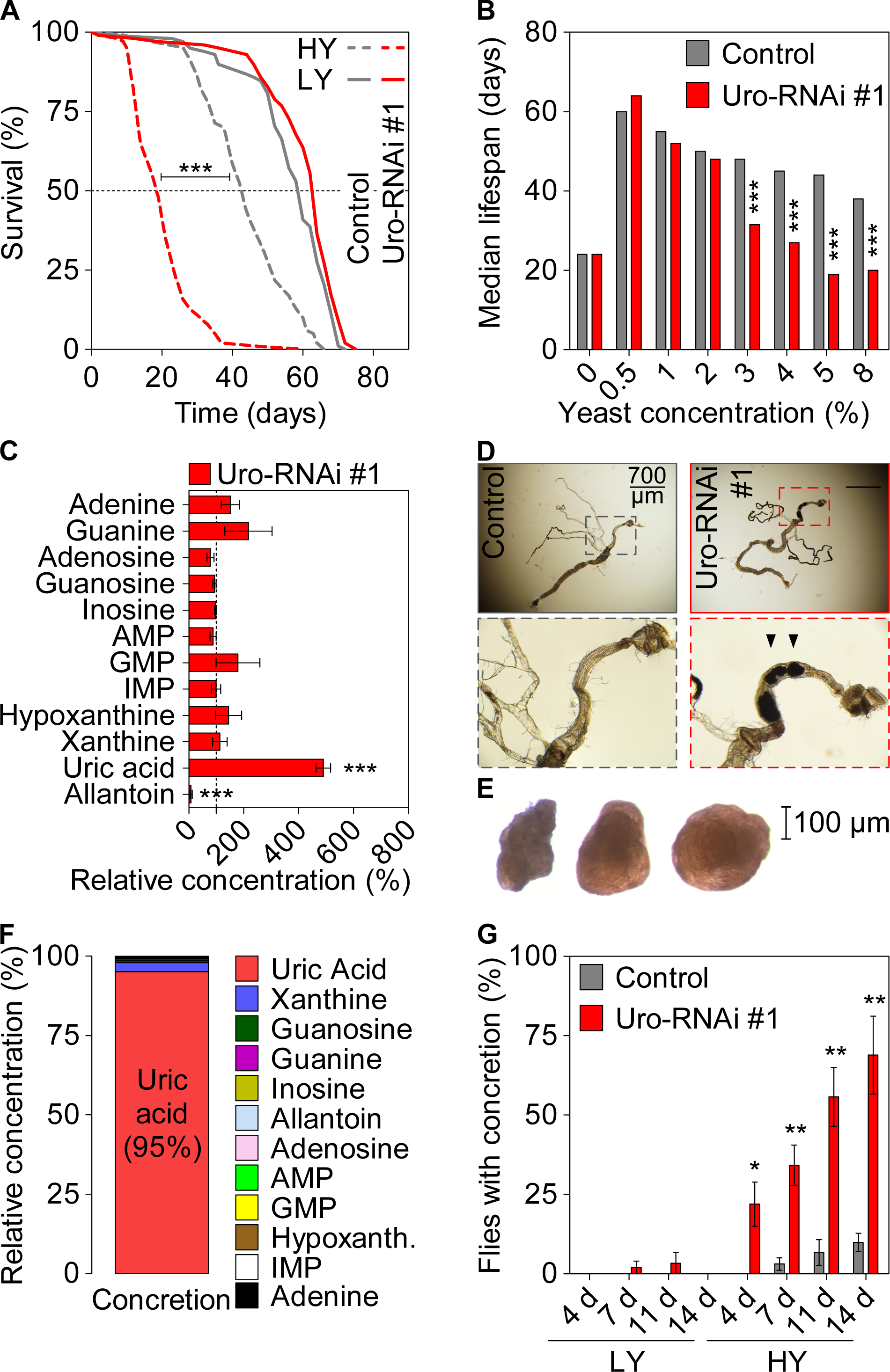
*Uro* knockdown attenuates lifespan and enhances UA concretion formation on a high yeast diet. (**A**) Survival curve of isogenic control and *Uro* knockdown flies (Uro-RNAi #1) on diets containing a low (LY; 0.5%) or high yeast extract content (HY; 5%). (**B**) Median lifespan of flies reared on diets containing increasing yeast extract contents. (**C**) Metabolite concentrations of fly homogenates from *Uro* knockdown flies relative to isogenic control flies reared on HY for 14 days. (**D**) Guts with attached Malpighian tubules of dissected flies. Arrowheads mark concretions of *Uro* knockdown flies. (**E**) Concretions extracted from the hindgut lumen of *Uro* knockdown flies. (**F**) Metabolomic analysis of extracted concretions. (**G**) Kinetic analysis of concretion formation determined by dissection. Supporting information is given in Figures S1, S2 as well as Movies S1 and S2.

Next, using *Uro* knockdown flies the metabolomic profiling verified the expected elevation of UA levels accompanied by a diminished allantoin production (Fig. 1C). Also, an unbiased micro-CT scan revealed radiopaque masses (putative ‘concretions’) specifically in the abdominal area of *Uro* knockdown flies, but not in the control population (Fig. S2A, Movie S1, S2). This observation was further substantiated by optical microscopy of dissected animals revealing the presence of concretions in the Malpighian tubule and hindgut of *Uro* knockdown flies (Fig. 1D, S2B). Due to the restricted expression pattern of the *Uro* gene to the excretory system both the Malpighian tubule and the hindgut represent the anatomical sites where UA is most likely expected to accumulate. Metabolomic analyses of isolated concretions confirmed UA as the predominant component, accounting for 95% of the metabolites measured (Fig. 1E, 1F, S2D). We also found enhanced UA aggregation (i.e., a high proportion of flies with UA concretions) with increasing age (Fig. 1G) and increasing dietary yeast content (Fig. S2C). Both effects match with the age- and diet-dependent occurrence of gout or uric acid kidney stones in humans (Doherty, 2009; Lieske et al., 2014). Consistent with the reduced lifespan phenotype, UA concretion formation was also observed when using other ubiquitous (*Da-GAL4*) and Malpighian tubule-specific (*c42-GAL4*, *Uro-GAL4*) *Uro* knockdown strains, but not in the neuronal-specific (*Elav-GAL4*) *Uro* knockdown strain (Fig. S2E).

### The dietary purine content mediates the lifespan and concretion phenotypes of *Uro* knockdown flies

To determine what component of the HY diet triggered increased UA production and associated phenotypes, we systematically supplemented the ‘benign’ low yeast diet (LY; 0.5% yeast content) with various basic nutrient groups: purines, pyrimidines, proteins, or sugar. We found that purines (metabolic precursors of UA), but not pyrimidines, proteins, or sugar led to a dose-dependent shortening of lifespan and concretion formation (Fig. 2A, 2B, S3). Purine homeostasis is controlled through orchestrated steps involving *de novo* synthesis, salvage, and degradation of purine intermediates (Fig. S4). The drug allopurinol, commonly used in the treatment of gout in humans, inhibits purine degradation by inhibiting xanthine dehydrogenase (XDH), an enzyme that converts hypo-/xanthine to urate. On the one hand, adding allopurinol to the HY diet rescued both the reduced lifespan and concretion formation in *Uro* knockdown flies (Fig. 2C, 2D, S5A, S5C), thus, supporting the validity of our translational model for UA based pathologies. On the other hand, inhibiting purine synthesis by adding methotrexate (inhibits folate biosynthesis) also rescued the phenotypes of *Uro* knockdown flies, suggesting an important role for purine *de novo* synthesis in UA homeostasis (Fig. 2E, 2F, S5B, S5C). Similarly, genetically knocking-down the targets inhibited by methotrexate or allopurinol, *dihydrofolate reductase* (*DHFR*) or *XDH*, in addition to *Uro*, significantly reduced UA concretion formation (Fig. S5D).

**Figure 2.**
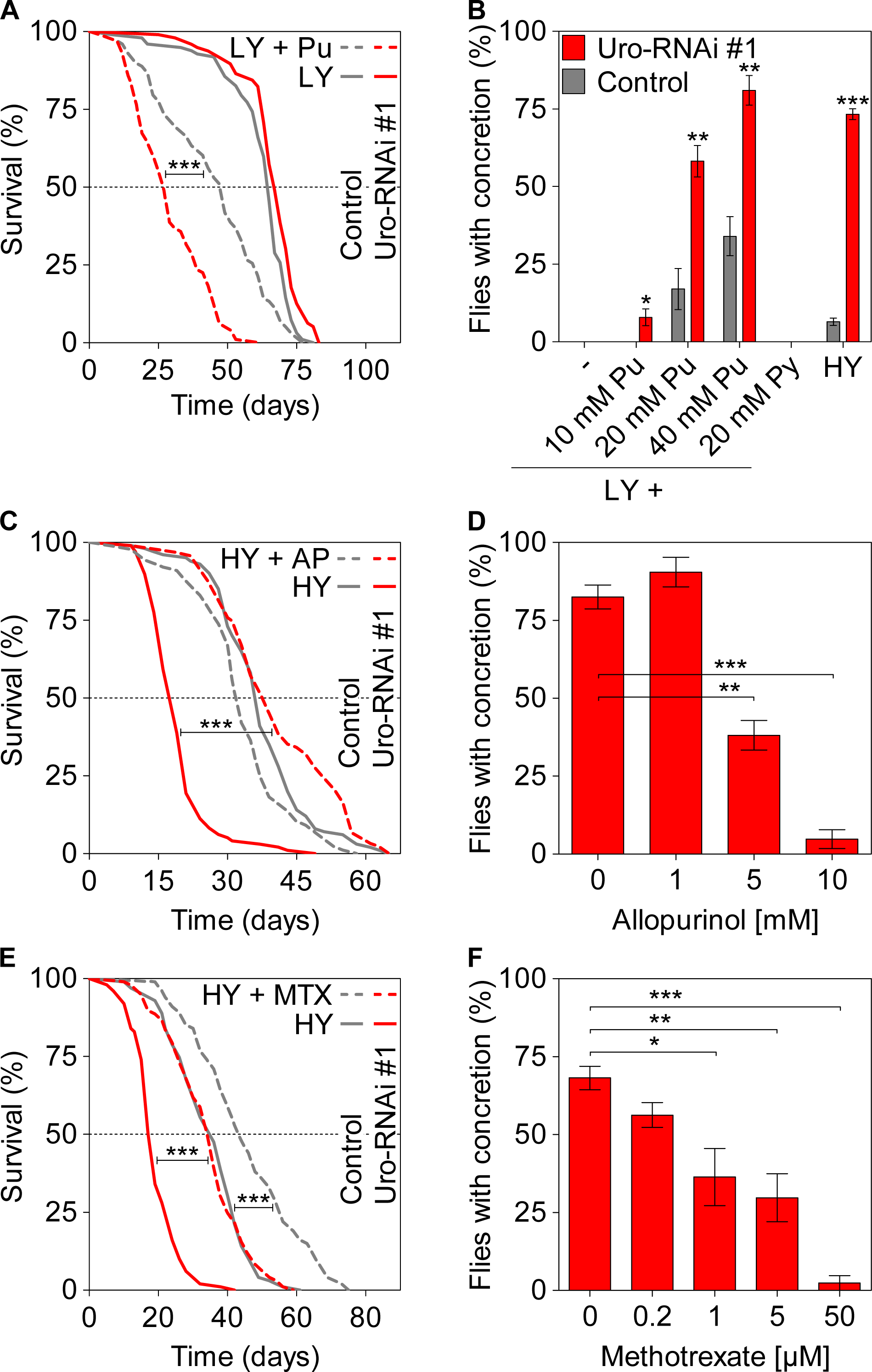
Dietary purine mediates lifespan attenuation and UA concretion formation. (**A**) Survival curve of control and *Uro* knockdown flies fed the LY or 40 mM purine supplemented (+ Pu) LY diet. (**B**) Concretion formation of flies reared on LY, purine (Pu) or pyrimidine (Py) supplemented LY, or HY diet for 14 days. (**C, E**) Survival curve of flies fed HY, 5 mM allopurinol (+ AP) or 5 µM methotrexate (+ MTX) supplemented HY diet. (**D, F**) Concretion formation of *Uro* knockdown flies reared for 14 days on HY without or with supplementation of allopurinol or methotrexate. Supplementary data is presented in Figures S3 – S5.

### The transcription factor FOXO dampens concretion formation by reducing UA levels and ROS formation

Next, with the *Drosophila* model at hand other potential genes involved in purine metabolism that might be targeted to control UA production were examined. This is of interest given, that (a) commonly prescribed medications such as allopurinol can result in serious adverse drug reactions and (b) common genetic variations identified to date explain only 6% of the variance encountered in serum UA levels, and consequently these genes might not represent ideal targets to lower the metabolic UA load (Hassan et al., 2011; Kinder et al., 2005; Reginato et al., 2012; Yang et al., 2010). To identify previously unrecognized regulators of UA metabolism, we examined the ILS pathway for two reasons. Firstly, ILS is a conserved signaling cascade activated when *Drosophila* is fed a HY diet (Britton et al., 2002; Broughton and Partridge, 2009; Russell and Kahn, 2007; Teleman, 2010). Secondly, a recent genome-wide association study identified polymorphisms in the human ILS gene *IGF1R* that were associated with serum UA concentration (Köttgen et al., 2013). Thus, we examined components of the ILS pathway for potential targets for future therapies to control UA production. Inhibiting the ILS pathway by inactivating either the insulin receptor (InR) or AKT kinase, or over-expressing *PTEN* (a negative regulator of human insulin signaling), resulted in a marked reduction in *Drosophila* concretion formation (Fig. 3A). Moreover, transgenic expression of *FOXO*, a downstream transcription factor inactivated by AKT kinase, reduced concretion formation (Fig. 3B) and UA production (Fig. 3C) in *Uro* knockdown flies.

**Figure 3.**
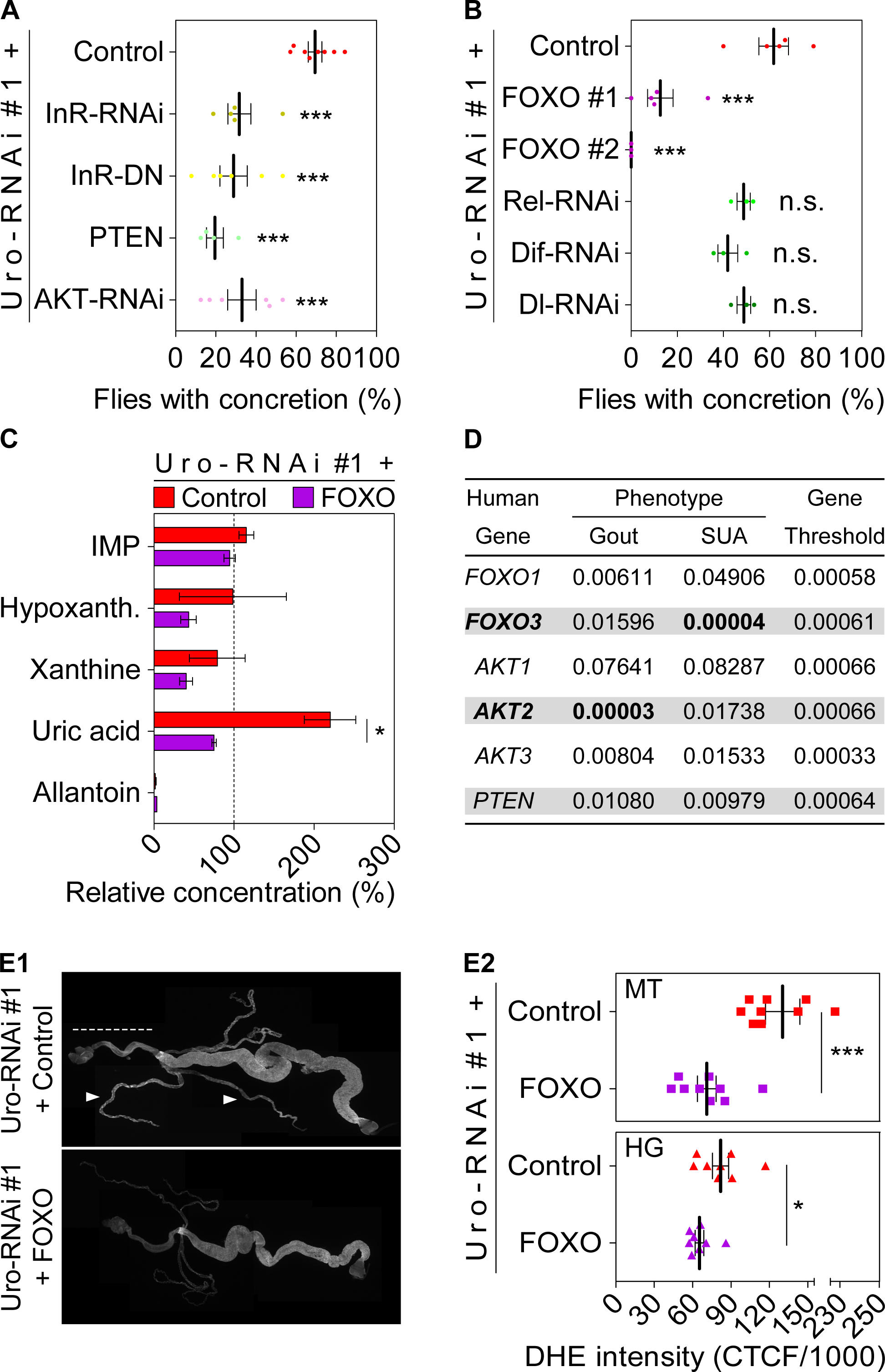
FOXO dampens concretion formation by reducing UA levels and ROS formation. (**A, B**) Concretion formation of flies expressing the *Uro* RNAi alone (Uro-RNAi #1 + Control) or in combination with a second transgene triggering either inhibition of insulin receptor (InR-RNAi, InR-DN (dominant negative)), AKT kinase (AKT-RNAi), or NFκB isoforms Relish, Dif, and Dorsal (Rel-, Dif-, Dl-RNAi), or triggering over-expression of PTEN (PTEN), or FOXO (FOXO #1, FOXO #2). (**C**) Metabolite concentrations of indicated fly genotypes. (**D**) P-values of SNPs within genes of the ILS network that significantly associated with gout or serum uric acid (SUA) levels in human subjects are indicated in bold. Significance of calculated p-values is set by the gene-specific threshold (last column). Effect sizes for significant SNPs are given in Tab. S2. (**E1**) Guts and Malpighian tubules from flies reared on HY diet were stained with DHE to visualize reactive oxygen intermediates. (**E2**) DHE staining intensity in the Malpighian tubules (MT) and the hindgut (HG) was quantified using the corrected total cell fluorescence (CTCF) of the respective organ area (McCloy et al., 2014). Supporting data is shown in Figure S6.

Yet, the translational potential of the above findings relies on there being a conserved mechanism between *Drosophila* and humans that relates the ILS pathway and UA levels. We assessed whether single nucleotide polymorphisms (SNPs) in different genes of the ILS pathway are associated with either UA levels or gout. Using data from humans (Tab. S1), we found that SNPs in the *AKT2* and *FOXO3* genes were associated with either UA levels or the incidence for gout (Fig. 3D, Tab. S2). These data support the idea that some constituents of the ILS signaling network, including the downstream transcription factor FoxO, play a critical and conserved role in affecting UA levels and associated pathology in both flies and humans.

### Lifespan attenuation and UA concretion formation of *Uro* knockdown flies are triggered by NOX mediated ROS production

We next addressed the underlying mechanism of the link between the ILS pathway and UA levels. UA has been proposed to play several physiological roles, depending on where it is acting. For instance, UA can act as an antioxidant in the blood, where it accounts for up to 60% of the antioxidative capacity (Maiuolo et al., 2016). In turn, in an intracellular setting such as in adipocytes, UA is considered a prooxidant and proinflammatory molecule (Imaram et al., 2010; Kanbay et al., 2016; Sautin and Johnson, 2008). Consistent with this, *Uro* knockdown flies showed elevated ROS production (prooxidative action) in the cells of the Malpighian tubule and hindgut accommodated by increased antimicrobial peptide expression (proinflammatory action). Both phenotypes could be reduced by *FOXO* over-expression (Fig. 3E1, 3E2, S6A-C). However, depleting the NFκB transcription factor paralogs (Relish, Dorsal, and Dif), whose function enhances expression of antimicrobial peptides, did not significantly diminish concretion formation (Fig. 3B). Therefore, it is likely that in the fly model the proinflammatory action secondary to *Uro* knockdown represents a downstream effect of UA, as seen in mammals (Lu et al., 2015; Lyngdoh et al., 2011), but does not itself cause any further elevation in UA levels.

The observed increase of ROS levels secondary to *Uro* knockdown (Fig. 3E2, S6A) could stem from two sources: (1) inefficient expression of ROS combating genes, or (2) an overabundance of ROS-producing enzymes. From the eleven oxidative stress-related genes whose expression was examined in the *Uro* knockdown flies only the ROS-producing *NADPH Oxidase* (*NOX*) showed significantly higher mRNA content (Fig. 4A, S6D). This increase in *NOX* expression was significantly abolished by *FOXO* over-expression (Fig. 4A). More directly, we also found that genetic ablation of *NOX* expression via RNAi countered both the short lifespan and concretion formation characteristically observed in *Uro* knockdown flies (Fig. 4B, 4C, S7). As with *FOXO* over-expression, knocking down *NOX* expression in *Uro* knockdown flies significantly reduced UA accumulation (Fig. 4D). We saw a similar result after pharmacologically inhibiting NOX activity and the associated ROS production using either apocynin (ACY; a NOX inhibitor) or vitamin C (Vit C; a ROS scavenger), respectively. Both treatments reversed the shortened lifespan and reduced concretion formation in a dose-dependent manner (Fig. 4E, 4F, S8A-C). While vitamin C seemed to extend the lifespan of both the control and *Uro* knockdown flies, the impact of apocynin on lifespan was specific for the urate oxidase depleted animals (Fig. 4E).

**Figure 4.**
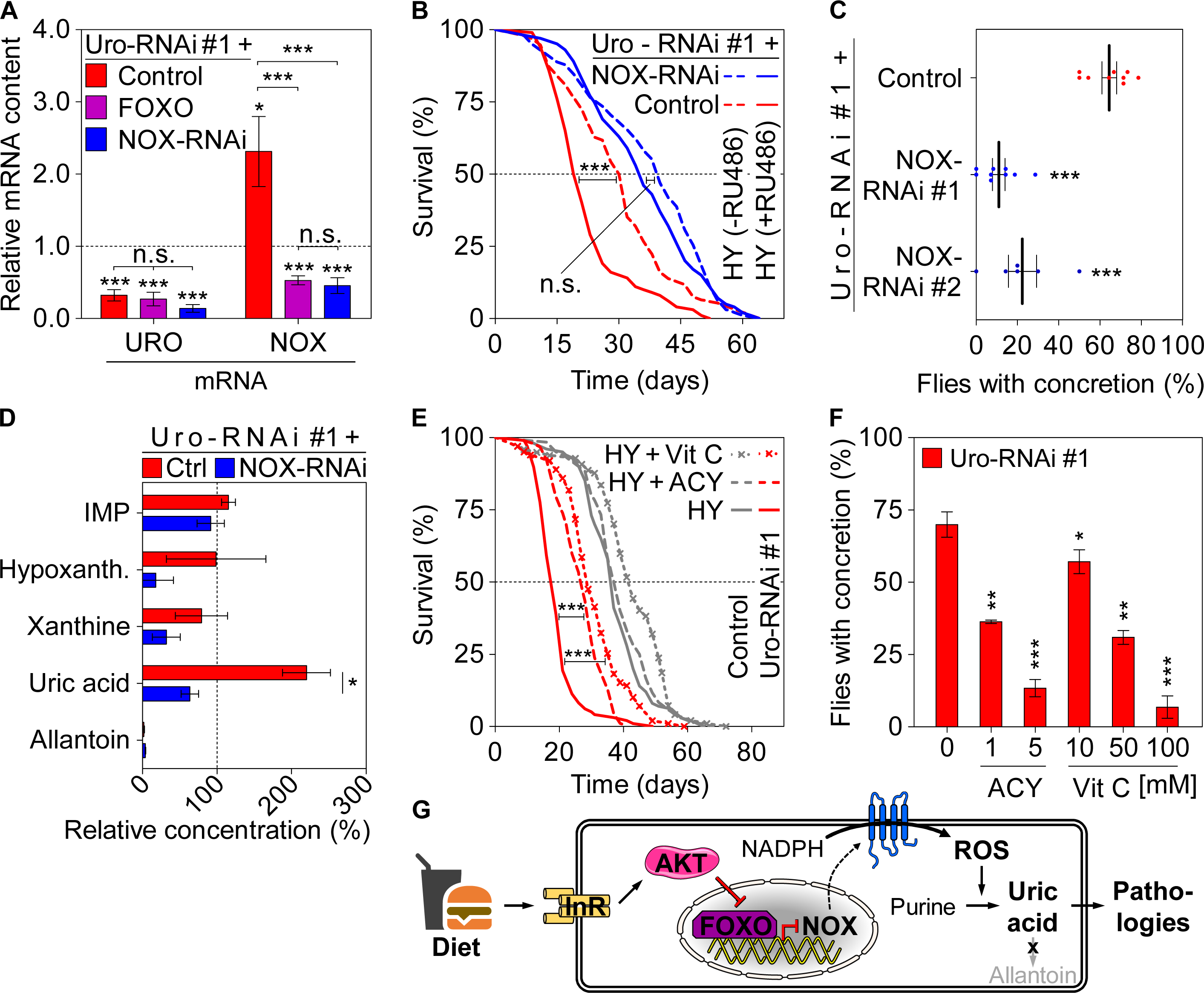
NOX mediated ROS production triggers lifespan attenuation and UA concretion formation on a high yeast diet. (**A**) *Uro* and NADPH oxidase (NOX) mRNA levels of flies expressing the Uro-RNAi alone (Uro-RNAi #1 + Control) or in combination with another transgene to over-express FOXO (FOXO) or inhibit NOX (NOX-RNAi). (**B**) Survival curve of flies carrying the Uro-RNAi transgene alone or in combination with a NOX-RNAi. Flies were fed a high yeast content (HY) diet without (-RU486) or with the transgene(s) activating RU486 ligand (+RU486) to assess the lifespan of the isogenic background and the corresponding single/double knockdown flies. (**C**, **D**) Concretion formation (**C**) and relative metabolite concentrations (**D**) of indicated fly genotypes. (**E**) Survival curve of flies fed HY without or with 5 mM apocynin (+ ACY) or 50 mM vitamin C (+ Vit C) supplementation. (**F**) Concretion formation of *Uro* knockdown flies reared for 14 days on HY without or with supplementation of apocynin or vitamin C. (**G**) Key players influencing UA levels and associated pathologies are summarized. Supporting data is shown in Figures S7 and S8.

Importantly, the transgenic *FOXO* or *NOX* manipulations did not alter the efficiency of *Uro* mRNA depletion, which argues for the direct impact of the *FOXO* and *NOX* gene products on UA level (Fig. 4A, the three bars to the left). Both alternatives over-expression of genes encoding for ROS-combating enzymes, such as cytosolic or mitochondrial superoxide dismutase (*SOD1*, *SOD2*), or treatment with the water-soluble vitamin E derivate Trolox failed to prevent UA concretion formation (Fig. S8C, S8D), and hence also argue for the direct involvement of NOX in modulating UA levels.

## Conclusion

Our studies argue the importance of purines as an important dietary component that limits lifespan. Generally, mutants in genes that influence lifespan upon dietary restriction either extend lifespan upon rich nutrient conditions while failing to extend lifespan upon dietary restriction conditions or attenuate the maximum lifespan upon dietary restriction. However, *Uro* mutants belong to a “novel” class of genes that limit lifespan only upon rich conditions but have little or no influence upon dietary restriction. We hypothesize that mutants that display such a phenotype encode a gene that amplifies the cellular damage that takes place under rich nutrient conditions compared to dietary restriction. We speculate that the increase of UA is deleterious and a lower UA level is partially responsible for the lifespan extension upon dietary restriction.

Our results also highlight the importance of ILS signaling, and its downstream effector NOX, as an important mediator of UA metabolism and related pathologies (Fig. 4G). Given the over-nutrition encountered in developed countries, the causal nexus uncovered here identifies potential new drug targets that could help regulate UA levels, ameliorating pathologies associated with hyperuricemia and extending human health-span.

## Acknowledgments

We thank the Bloomington Stock Center and Vienna Drosophila RNAi Center for providing the fly strains. We also thank members of Kapahi lab for discussion and suggestions. This work was funded by grants from the American Federation of Aging Research and Hillblom foundations and NIH (R01AG038688 & RO1AG045835) granted to Pankaj Kapahi. Development of the RPGEH was supported by The Robert Wood Johnson Foundation, the Wayne and Gladys Valley Foundation, The Ellison Medical Foundation, KPNC, and the Kaiser Permanente National and Regional Community Benefit Programs. Work and personnel for this project were funded by NIH grant R01DK103729. Deanna Brackman was additionally funded by NIH Training Grant T32 GM717537.

## Author Contributions

SL designed, executed and analyzed the *Drosophil*a experiments with support from JNB, KAW and TH. NB performed the metabolomic analyses including sample preparation with support from AS. MW did the DHE staining and analysis. LC and SH supervised the µCT imaging. DJB and KG performed the human CGS. SL, DWK, AK, MLS, TC, and PK supervised experiments and wrote parts of the manuscript. All authors read and contributed to the final text version.

## Material and Methods

### Contact for reagent and resource sharing

Further information and requests for resources and reagents should be directed to and will be fulfilled by the Lead Contact, Pankaj Kapahi.

## Experimental model and subject details

### Fly strains used in this study

Fly strains purchased from Bloomington Drosophila Stock Center are indicated by the prefix “b”, strains from FlyORF, Zurich by “F”, strains from NIG-Fly, Mishima by “n”, and strains from VDRC, Vienna by “v”.

*GAL4* driver lines: daughterless gene switch (kindly provided by Linda Patridge), daughterless (b55851), c42 (b30835), Uro (b44416), Elav (b458)

*UAS* responder lines: Uro-RNAi #1 (n7171-R1), Uro-RNAi #2 (n7171-R2), InR-RNAi (b31594), InR-DN (b8252), PTEN (kindly provided by Tiang Xu), AKT-RNAi (b31701), FOXO #1(b9575), FOXO #2 (F000143), Rel-RNAi (b28943), Dif-RNAi (b29514), Dl-RNAi (b32934), NOX-RNAi #1 (b32902), NOX-RNAi #2 (b32433), Gal4-RNAi (b35784), PRPS-RNAi (b35619), DHFR-RNAi (b35015), XDH-RNAi (v25175), SOD1 #1 (b24750), SOD1 #2 (b33605), SOD2 (b24494)

Other lines: Control A (W1118), Control B (b35785), Control C (b36303), Control D (v60100). We generated a recombinant fly line carrying both genetic elements (+/+; DaGS::Uro-RNAi #1/ DaGS::Uro-RNAi #1; +/+) on chromosome two. This line was crossed to *UAS* responder lines to introduce an additional transgenic element or one of the control lines (Fig. 3, A-C and E, Fig. 4, A-D, and Figs. S5D, S6C, S7, S8D).

### Fly husbandry, dietary and pharmacological manipulations

All fly lines were maintained on standard fly yeast extract medium containing 1.5% yeast, 5% sucrose, 0.46% agar, 8.5% of corn meal, and 1% acid mix (a 1:1 mix of 10% propionic acid and 83.6% orthophosphoric acid) prepared in distilled water. To prepare the media, cornmeal (85 g), sucrose (50 g), active dry yeast (16 g, "Saf-instant") and agar (4.6 g) were mixed in a liter of water and brought to boil under constant stirring. Once cooled down to 60°C 10 ml of acid mix was added to the media. The media were then poured in vials (~10 ml/ vial) or bottles (50 ml/bottle) and allowed to cool down before storing at 4°C for later usage. These vials or bottles were then seeded with some live yeast just before the flies are transferred and used for maintenance of lab stocks, collection of virgins, or setting up crosses.

Experimental low and high yeast diets varied only with regard to the yeast type and content ranging from 0% to 8%. Diets declared as LY (low yeast content) and HY (high yeast content) contained 0.5% and 5% Baker’s yeast extract (#212750 BactoTM Yeast Extract, B.D. Diagnostic Systems, Sparks, MD), respectively. If not stated otherwise pharmacological treatments or dietary supplementations were mixed with the other ingredients during preparation of the food. As for the standard medium, the mix was brought to a boil under constant stirring and allowed to cool down to 60°C before adding the acid mix. For the induction of transgenic elements via the *GAL4-UAS* system when using a gene switch driver 200 μM RU486 (Mifepristone, Sigma) in 100% ethanol was added to the food while the isogenic controls received vehicle treatment. The media was then poured in vials (~5 ml/vial) and allowed to cool down before storing at 4°C for later usage.

The dietary supplementations and drug treatments used were as follows:

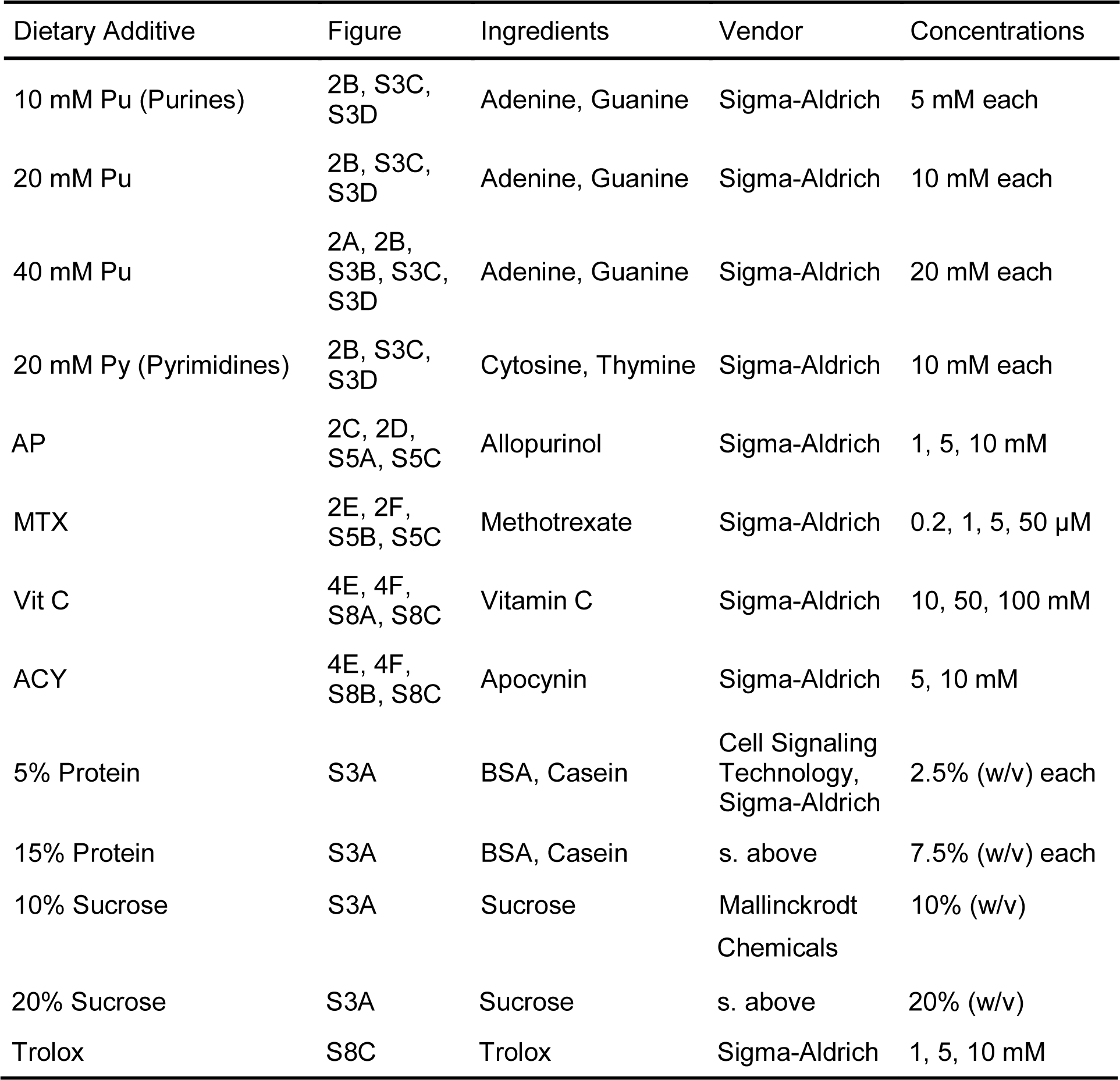

### Fly Rearing

Genetic crosses were obtained by pairing 15 young virgin females (carrying the *GAL4* driver construct) with 4 young male flies (carrying the *UAS* responder construct) in new stock bottles. Flies were kept in the stock bottles for five days, after which the parents were removed, and the larvae could develop in a temperature (25°C) and humidity (60%) controlled designated fly room with a 12-hour light / dark cycle. Newly eclosed flies could mate for 2 - 3 days, to complete development post-eclosion, before they were grouped under light CO_2_ anesthesia. Sorted females were then transferred to the appropriate media vials for subsequent analyses.

## Method details

### Lifespan Analysis

Flies developed on standard fly 1.5% yeast extract medium were transferred to the necessary diet within 72 hours after eclosion. For survivorship analysis four to six media containing vials with 25 mated females were kept in a temperature (25°C) and humidity (60%) controlled designated fly room with a 12-hour light / dark cycle. Every other day flies were transferred to fresh food and fly survival was scored by counting the number of dead flies. The significance of change was determined using the log-rank test.

### Dissection and scoring of concretion formation

For dissection and imaging assays adult female flies were dissected after 4, 7, 11, or 14 days. The excretory system, in particular the hindgut and Malpighian tubule area was checked and imaged for the presence of ectopic biomineralization. Flies were anesthetized by CO_2_ on standard flypads (Cat # 59–108, Genesee Scientific, San Diego, CA) and dissected under a dissecting light microscope (SZ61, Olympus, Center Valley, PA) on a culture dish in droplets of distilled water utilizing fine forceps (Roboz ceramic, Gaithersburg, MD). The excretory system was imaged utilizing a Zeiss Stereo Microscope with external light source. While transparent posterior midguts and hindguts devoid of any sign of deposit formation were scored as negative, those with pale yellow/orange hard concretions were scored as deposit carrying specimens. The relative level of deposit formation was quantified by this binary approach. Per condition at least 15 - 20 flies were dissected and the number of flies with deposits was divided by the total number of flies dissected. Each experiment was repeated for a minimum of three independent, biological replicates.

### Imaging of ectopic biomineral

Images of dissected guts and Malpighian tubules were captured with the Olympus SZX12 microscope. Isolated deposits extracted from the hindgut area were imaged with the Olympus BX51 microscope (Olympus Scientific Solutions Americas Corp. Waltham, MA). For the micro-computer tomography (µCT) imaging fly specimens were scanned in 50% ethanol using a micro-XCT unit at 4X magnification (4.5 µm/voxel) with 1200 image projections. Scanning was performed with a 3 second exposure, source power of 40 W, 200 µA current, source distance of ~30 mm, detector distance of ~14 mm, camera binning 2, and an angle sweep from −93° to 93°. Post-3D reconstruction, the acquired data was further processed and analyzed with Avizo software (Version 9.3.0; FEI, Hillsboro, Oregon).

### Collection of deposits for mass spectrometry-based metabolomics

Flies were dissected in 50% methanol/water mix and the deposits from the hindgut area were isolated using fine forceps. Deposits of 25 flies carrying concretions were collected in an Eppendorf tube filled with 500 µl 50% methanol/water. The material was sedimented by centrifugation for 15 s with 5,000 rpm and the supernatant was aspirated. After two further wash steps with 500 µl 50% methanol/water deposits were frozen in liquid nitrogen and stored at - 80°C until further processing.

### High-performance liquid chromatography (HPLC) mass spectrometry (MS)

High-performance liquid chromatography (HPLC) was performed using an Agilent 1260 UHPLC system and connected to a Phenomenex Luna NH2 column (2 × 100 mm, 3 μm, 100 Å) and a SecurityGuard™ NH2 guard column 4 × 2 mm ID. Mass spectrometry (MS) was performed using a 5500 Triple-Quadrupole LC-MS/MS mass spectrometer from Sciex fitted with a Turbo VTM ion source. Sciex’s Analyst^®^ v1.6.1 (MacLean et al., 2010) was used for all forms of data acquisition, development of HPLC method, and optimization of analyte-specific MRM (multiple reaction monitoring) transitions. Skyline^®^ v4.1 was used for LC-MS/MS data analysis. For whole fly analysis, 5 flies (in sextuplicates) per condition were flash-frozen over liquid nitrogen and subsequently homogenized ultrasonically using a Fisher Scientific’s 550 Sonic Dismembrator with 50 μl of an 8:2 mixture of methanol/water (v/v), containing 2.5 µM of 2-chloroadenosine as internal standard. Three 20 s pulses at amplitude setting 3 of the instrument (on ice) were sufficient to completely homogenize fly specimens. The homogenates were then vortexed for 5 times over a period of ~30 min (each 1 min long). Subsequently, the samples were centrifuged at 10,000 rpm for 10 min, the supernatant was filtered, and 3 μl of each was injected for HPLC-MRM analysis (*vide infra*) without any additional processing. For the analysis of fly deposits, samples were collected as described above and prepared for HPLC-MRM analysis (*vide infra*) as described above for whole flies.

Optimization of analyte-specific MRM transitions, such as determination of suitable precursor and product ions and optimal MS parameters for each transition (Q1, precursor → Q3, product) were achieved by isocratic flow injection of the 1-10 μM solution (final) for each standard, diluted in 80% methanol. The most intense transition was used as quantifier, whereas one or more additional transitions were used as qualifier for each compound (Tab. S3). A final standard mixture of all compounds at 5 µM (containing the internal standard 2-chloroadenosine at 2.5 µM), were prepared prior to analysis and injected at the onset of each biological sample set. Based on previous reports (Yuan et al., 2012), the following HPLC program was developed: a solvent gradient of 20 mM ammmonium acetate + 20 mM ammonium hydroxide (pH=~9.5) + 5% acetonitrile (aqueous) – acetonitrile (organic) was used with 0.4 ml/min flow rate, starting with an acetonitrile content of 85% for 1 min, which was decreased to 30% over 3 min and then to 0% over 7 min and held at 0% for 2 min. The HPLC column was subsequently reconstituted to its initial condition (acetonitrile content of 85%) over the next 1 min and re-equilibrated for 7 min. Metabolome ex-tracts from whole flies or deposits were analyzed by HPLC-MRM with positive/negative switching of source ion modes (Yuan et al., 2012). Source conditions were as follows: curtain gas (CUR) 20, collision gas (CAD) 7, ion source gas 1 (GS1) 30, ion source gas 2 (GS2) 30, ionspray voltage (IS) ±4500 V, and source temperature (TEM) 450°C. Quantification was based on integration of analyte-specific peaks obtained from HPLC-MRM runs.

### Dihydroethidium (DHE) staining for ROS measurements

Intact guts with adhering Malpighian tubules were dissected from female flies in a droplet of PBS (137 mM NaCl, 2.7 mM KCl, 10 mM Phosphate, pH 7.4) on ice and subsequently stained for 5 min with 45 µM DHE (Thermo Fisher Scientific) dissolved in DMSO (Sigma Aldrich). After three 10 min wash steps with PBS the sample was fixed for 45 min using 4% formaldehyde (Sigma Aldrich) in PBS followed by another three 10 min wash steps. After a 15 min staining with 1 µg/ml DAPI (Thermo Fisher Scientific) dissolved in PBS the specimen was mounted onto a microscope slide using Mowiol (Sigma Aldrich). Confocal images were collected using a Zeiss LSM700 confocal system using an Excitation/Emission (nm) of 518/605. DHE intensity was quantified using Image J.

### Total RNA and cDNA preparation

Total RNA was isolated from 5-10 females using the Quick-RNA™ MiniPrep Kit (Zymo Research) at room temperature. In brief, flies were anesthetized with CO_2_ before homogenization using a Kontes Microtube Pellet Pestle Rods with Motor in an Eppendorf tube containing 200 μl RNA Lysis Buffer. After adding another 400 µl RNA Lysis Buffer the homogenate was centrifuged for 1 min with 12.000 rcf and 400 µl supernatant was transferred to the spin-away filter placed over a fresh collection tube for elimination of genomic DNA. 400 µl of 95 % ethanol was added to the flow through and mix. Transfer mixture to Zymo-Spin IIICG column in a collection tube and centrifuge for 1 min with 12,000 rcf. After adding 400 µl RNA Prep Buffer and a 30 sec centrifugation with 12,000 rcf two more wash steps with 700 µl and 400 µl RNA Wash Buffer follow. To ensure complete removal of the RNA Wash Buffer the last centrifugation is carried for 2 min with 12,000 rcf. The Zymo-Spin IIICG column is placed in a collection tube and bound RNA is eluted with 30 µl DNAse/RNAse-free water in a collection tube by centrifugation for 30 sec with 12,000 rcf. Quantity and quality of isolated RNA was determined using the NanoDrop 1000 Spectrophotometer (Thermo Scientific).

1 µg of total RNA in a volume of 16 µl was used per sample and cDNA was synthesized using 4 µl iScript Reverse Transcription Supermix for RT-qPCR (Bio-Rad) according to the manufacturer’s protocol. The RT-PCR reaction protocol included a priming step (5 min at 25°C) followed by the reverse transcription (30 min at 42°C) and inactivation of the reaction (5 min at 85°C). If necessary, cDNA was stored at −20°C before the quantitative real-time PCR.

### Quantitative real-time PCR (qPCR)

Using 3 µl of the 1:10 diluted cDNA as a template, 2 µl of a primer pair (500 nM each) and 5 µl SensiFAST SYBR No-ROX Kit (BIOLINE) the qPCR was performed using the Light Cycler 480 Real-Time PCR System (Roche Applied Science). After pre-incubating the sample (95°C for 2 min) to denature DNA forty PCR cycles of denaturing (95°C for 5 s, ramp rate 4.8°C/s), as well as annealing and extension (60°C for 20 s, ramp rate 2.5°C/s) followed. The specificity of amplicons was verified with a subsequent melting curve analysis. The data was analyzed by means of the ΔΔCt method and the values were normalized using β-tubulin as an internal control. The following primer pairs were used in this study:

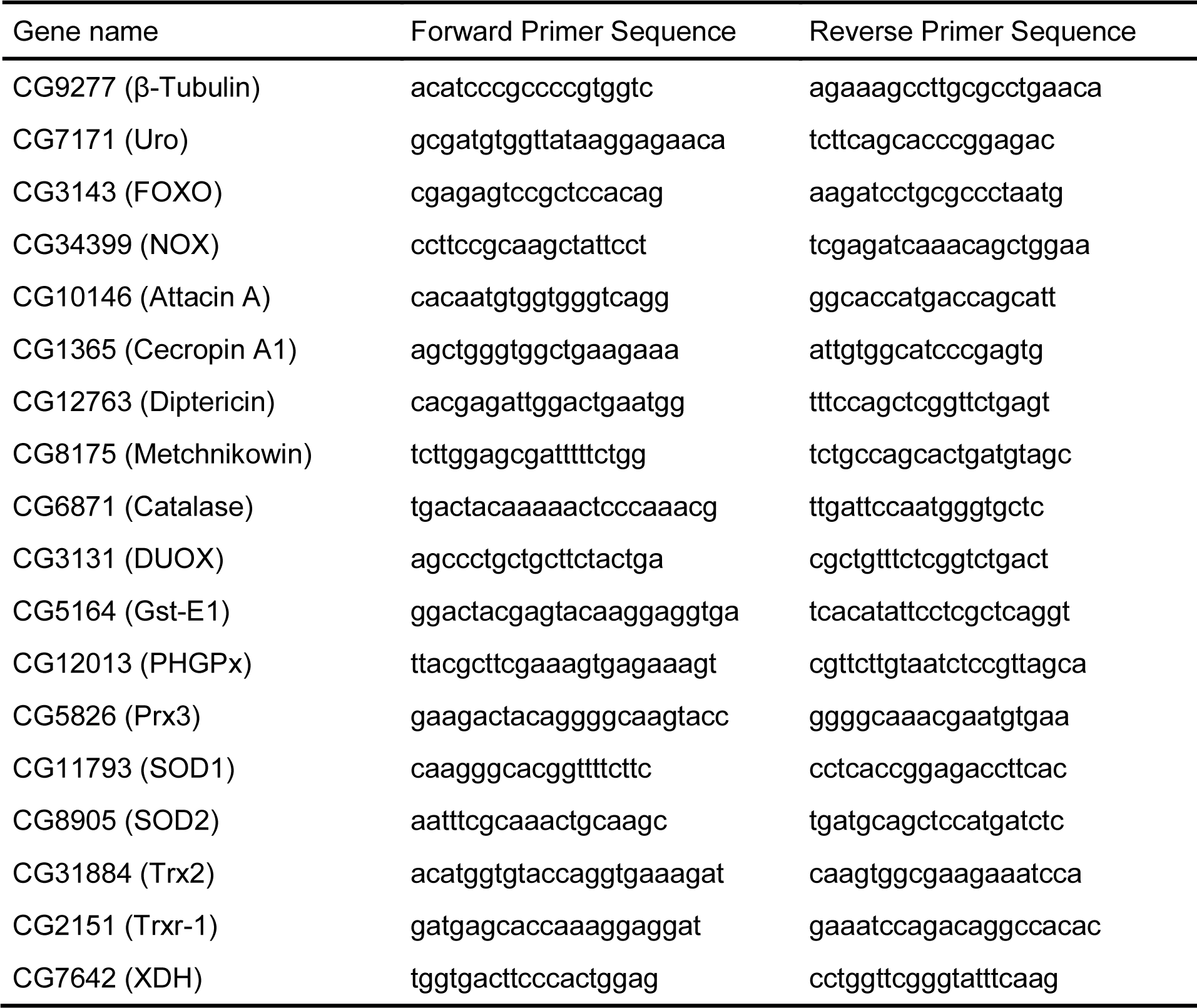

### Human candidate-gene study (CGS)

Data from the Kaiser Permanente Epidemiology Research on Adult Health and Aging (GERA) Cohort, a subset of the Research Program on Genes, Environment and Health (RPGEH), were gathered for analysis. Detailed methods on the genotyping, imputation, and ancestry calculations can be found in (Wen et al., 2015). The study population consisted of 80,795 adult subjects of European ancestry. The average age of the cohort was approximately 63 years old at the time of sample collection for genotyping. Two phenotypes were used for this study: (1) Gout, defined as two ICD-9 codes for gout in the electronic health record; (2) mean serum UA levels. Association analyses were conducted using linear or logistic regression models in PLINK v1.90 adjusting for population structure using the top 10 principal components. In the model, concomitant diuretics and gender were included as covariates because of their strong correlation with all phenotypes. Maximum body mass index (BMI) correlated strongly with UA levels, and was therefore also included as a covariate in all analyses.

SNPs within +/-2kb of ILS genes (*FOXO1*, *FOXO3*, *AKT1*, *AKT2*, *AKT3*, *PTEN*) were analyzed. Lists of SNPs within genes were analyzed using NIH LDLink SNPclip tool (Machiela and Chanock, 2015), where SNPs in high linkage disequilibrium (R^2^>0.5) were pruned. Gene-specific p-value thresholds were set as 0.05/(2 phenotypes*number of SNPs in each gene post-pruning). As a quality control step, top associated SNPs were checked for extreme HWE departures in our samples.

R v3.4.0 was used to create phenotypes and analyze associations between risk factors and phenotypes.

### Quantification and statistical analysis

Bar graphs and scatter blots represent the mean ± standard error of the mean (SEM), when indicated. Lifespan curves, and other graphs were visualized using GraphPad Prism 5 software. For statistical comparisons between the control and single treatment group a two-tailed student’s t-test was used; except for lifespan analysis where the log-rank test was performed. When multiple conditions were compared against one control group ANOVA was used. Significant differences are indicated according to the p-value by asterisks with p < 0.001 (***) < 0.01 (**) < 0.05 (*). Non significant differences are indicated by n.s.

